# Apoptosis uncovers human evolutionary heritage sequestered in non-coding DNA

**DOI:** 10.1101/2025.11.13.688179

**Authors:** Ruchi Joshi, Anushka Roy, Relestina Lopes, Laxmi Kata, Rutuja Selukar, Mrunmayi Markam, Subhangi Banerji, Gorantla V Raghuram, Snehal Shabrish, Indraneel Mittra

**Affiliations:** Translational Research Laboratory, Advanced Centre for Treatment, Research and Education in Cancer, Tata Memorial Centre, Kharghar, Navi Mumbai 410210, India; Homi Bhabha National Institute, Anushakti Nagar, Mumbai 400094, India

## Abstract

Cell-free chromatin particles (cfChPs), primarily derived from non-coding DNA (ncDNA) and released from apoptotic cells into the bloodstream, can be horizontally transferred into living cells. However, their potential role in generating ncDNA within recipient cells remains unknown. We previously reported that cfChP- sized fragments generated upon sonicating high-molecular-weight DNA can indiscriminately transmit themselves into foreign cells across species and kingdom boundaries. We hypothesized that cfChP-like DNA–protein complexes generated following organismal death have been exchanged among species and have progressively accumulated to form a dense patchwork that now constitutes the ncDNA, and that apoptosis may dissociate these complexes, rendering them amenable to detection. To test this hypothesis, we used species-specific DNA fluorescent in situ hybridisation probes and antibodies against archaea, eubacteria, plants, algae, fungi, protozoa, Drosophila, fish, chicken, mouse, rat, pig, dog, and monkey on intact and apoptotic human cells. We detected no reactivity with intact cells but strong reactivity with the ncDNA component of apoptotic cells. These findings indicate that ncDNA represents a dense conglomeration of DNA–protein complexes derived from different species that become detectable following apoptosis. This suggests that human evolutionary heritage is conserved within ncDNA formed through progressive accumulation of genetic fragments transferred horizontally following organismal death.

## Main

The genomes of complex multicellular eukaryotes are primarily composed of non-protein-coding DNA (ncDNA). Although a fraction of these genomes has important regulatory functions^1^, the reason why 99% of the human genome should comprise ncDNA remains unresolved.

We previously reported that circulating cell-free chromatin particles (cfChPs), primarily composed of ncDNA fragments released from apoptotic cells, undergo horizontal transfer into living cells, where they become detectable using fluorescent in situ hybridization (FISH)^2^. These findings suggested that DNA sequences, typically undetectable within the ncDNA of intact cells, can be visualised following apoptosis, albeit within foreign host cells. We also demonstrated that ncDNA contained within cfChPs harbours an intrinsic protein synthesis machinery, including RNA polymerase, ribosomal RNA, and ribosomal proteins, enabling them to autonomously synthesize various proteins that are typically produced by genomic DNA. These proteins can be detected using immunofluorescence (IF) within host cells with the aid of appropriate antibodies.

In a related study, we reported that mechanical shearing of high-molecular-weight DNA via sonication results in the generation of cfChP-like small DNA–protein complexes, which can readily enter foreign cells without being restricted by species and kingdom boundaries³. Based on this finding, we hypothesized that cfChP-like DNA–protein complexes generated upon organismal death, especially during evolutionary periods of acute environmental stress^4–6^, were exchanged among species and progressively accumulated to form the dense patchwork that now constitutes the ncDNA. We further hypothesized that apoptosis may dissociate the constituents of the compacted ncDNA, making them amenable to detection by FISH and IF^2^.

To test this hypothesis, we used species-specific FISH probes to detect ncDNA sequences unique to various taxa, from archaea to monkeys, within apoptotic human cells, using intact cells as negative controls. Since cfChPs, like DNA– protein complexes, can autonomously synthesize proteins^2^, we also tested whether multiple species-specific antibodies react with intact and apoptotic cells. Before we began our experiments, we confirmed that our FISH probes and antibodies were strictly species-specific and exhibited no cross-reactivity (Supplementary Fig. 1).

We believe that our findings reported herein will provide insights into the evolutionary origin and composition of ncDNA, clarifying whether interspecies exchange of DNA–protein complexes has contributed to the progressive expansion of the non-coding genome throughout evolution. The latter would be consistent with emerging reports that evolutionarily advanced species possess progressively higher fractions of ncDNA^7–9^.

### Hypoxia-induced apoptosis

We first investigated whether apoptosis induced by hypoxia or occurring naturally in human cells and tissues reacts with species-specific DNA FISH probes and antibodies, whereas intact living cells do not. Accordingly, we subjected A375 human melanoma cells to hypoxia (1% oxygen). We exposed the flow-sorted Annexin V-positive apoptotic cells and the Annexin V-negative non-apoptotic (living) cells to nine different species-specific FISH probes and antibodies. Cells or tissues from each of these species acted as positive controls (Figs. 1a and 1b). We observed that the FISH probes and antibodies did not react with intact A375 cells but reacted strongly with apoptotic A375 cells and positive-control cells/tissues. We validated these findings in A375 cells by performing similar experiments using four additional human cell lines, HEK-293 (embryonic kidney), HeLa (cervical cancer), HEKa (epidermal keratinocytes), and HADF (adult dermal fibroblasts), with results identical to those obtained in A375 cells (Supplementary Fig. 2).

**Fig. 1:**
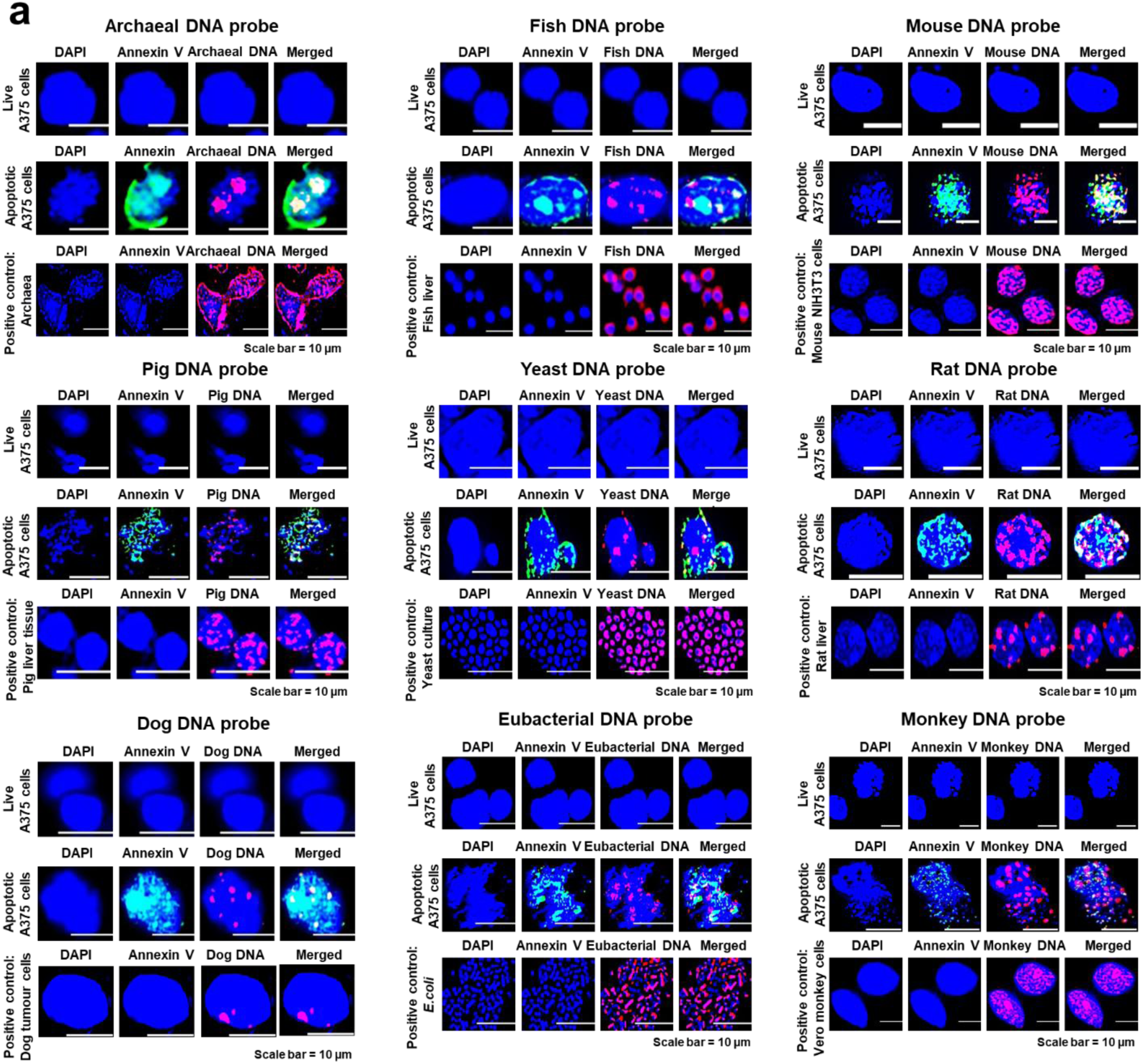
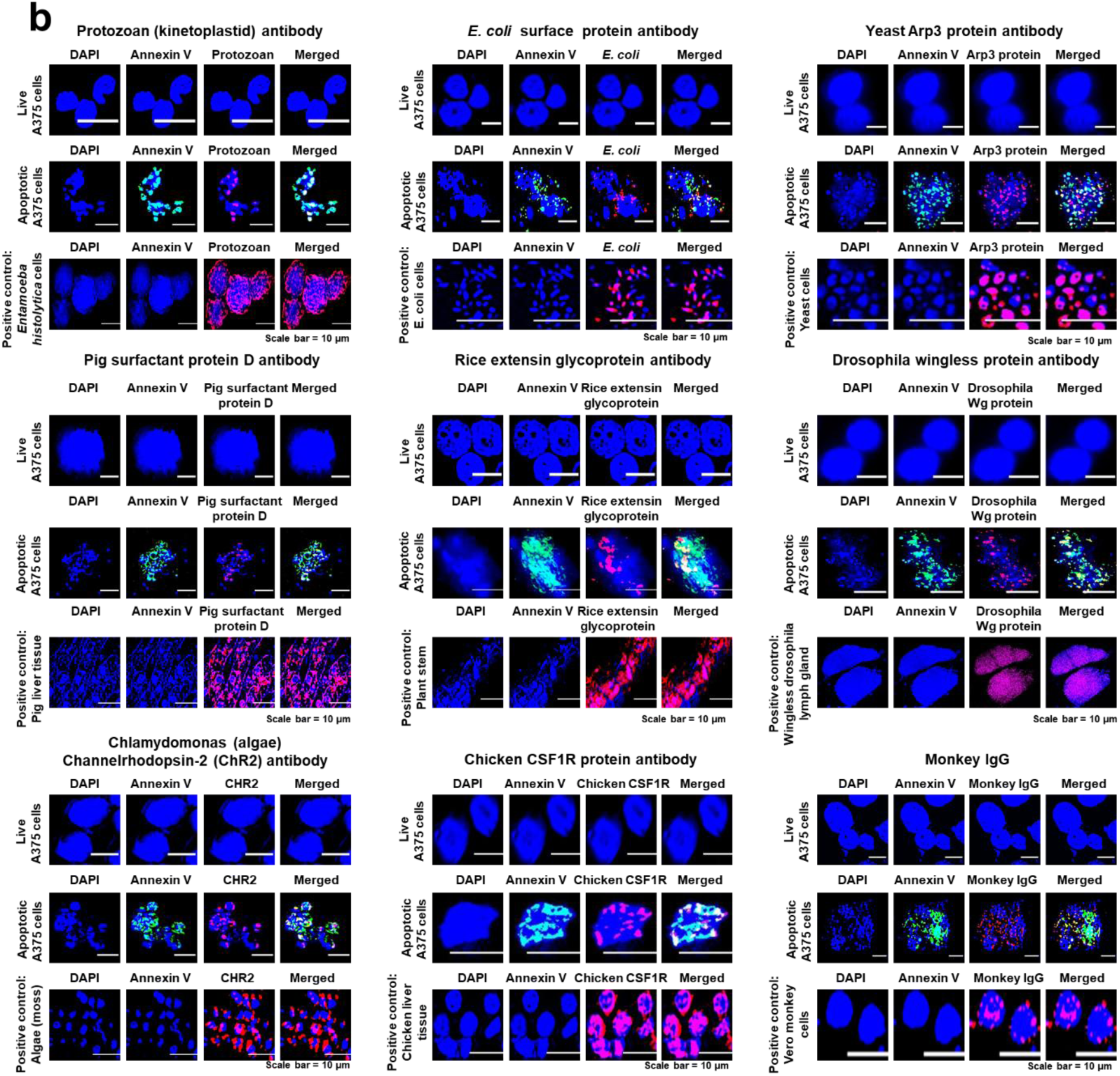
Detection of species-specific nucleic acid and protein markers in apoptotic human melanoma cells. **a.** Live A375 cells and the corresponding flow-sorted Annexin V-positive apoptotic cells were simultaneously probed with species- specific DNA FISH probes. Apoptotic cells (middle row in each panel) but not live cells (top row in each panel) reacted with the FISH probes. The bottom row in each panel represents positive-control cells or tissues of the respective species. The dog FISH probe used was specific for the X (magenta) and Y (green) chromosomes. As A375 cells originated from a female patient with malignant melanoma, only the magenta X chromosome is visible in the apoptotic A375 cells. Similarly, green signals (Y chromosome) are not observed in the positive-control cells, as they were derived from a female dog tumour. **b.** Live A375 cells and the corresponding flow-sorted Annexin V-positive apoptotic cells were simultaneously probed with species-specific antibodies. Apoptotic cells (middle row in each panel) but not live cells (top row in each panel) reacted with the antibodies. The bottom row in each panel represents positive-control cells/tissues of the respective species. All experiments presented in Fig. 1 were performed twice by two different investigators.

### Spontaneously occurring apoptosis

We next investigated the reactivity of cells that had spontaneously undergone apoptosis under standard culture conditions or in normal human tissues. We examined three human cell lines, namely, MDA-MB-231 (breast cancer), MRC-5 (embryonic lung fibroblast), and A375 (malignant melanoma), as well as three normal human tissues (brain, liver, and kidney). We identified spontaneously occurring apoptotic cells among the viable cells via immunostaining with an antibody against Annexin V. We observed that FISH probes against rat, archaea, fungal, fish, bacterial, and mouse DNA reacted strongly with Annexin V-positive apoptotic cells but not with Annexin V-negative viable cells (Fig. 2a, top row in each panel). Annexin V-positive apoptotic cells also reacted with antibodies against protozoan kinetoplastid, rat RTA1 protein, rice extensin glycoprotein, mouse MHC- II protein, E. coli O15 protein, and chicken CSF1R protein. In contrast, Annexin V-negative viable cells were non-reactive (Fig. 2a, bottom row in each panel).

**Fig. 2:**
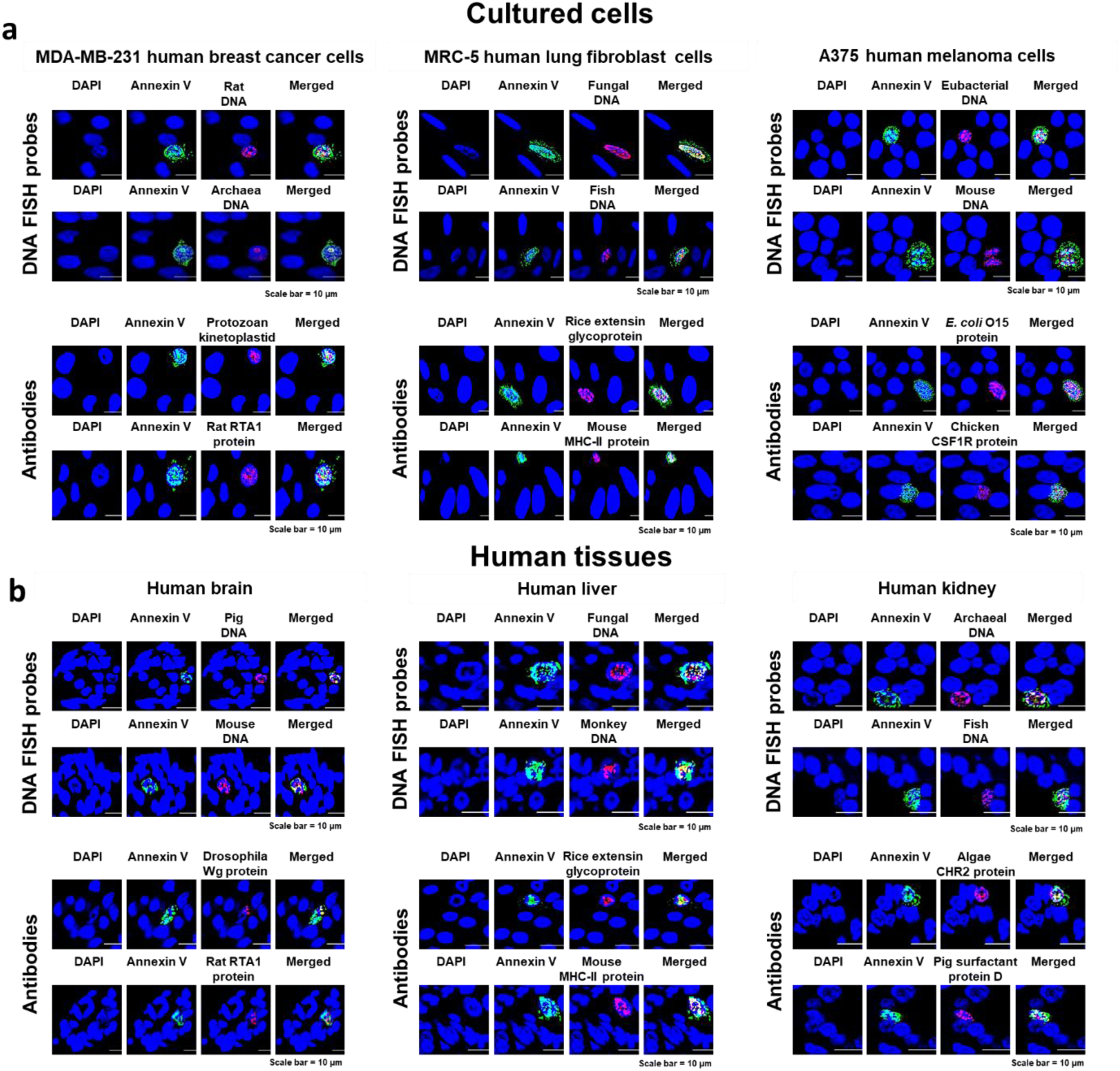
Spontaneously occurring apoptotic cells, but not viable cells, reveal phylogenetic diversity. **a.** MDA-MB-231 human breast cancer, MRC-5 human embryonic lung, and A375 human melanoma were simultaneously stained with an antibody against Annexin V and a panel of species-specific DNA FISH probes and antibodies. Annexin V-positive apoptotic cells, but not Annexin V-negative viable cells, exhibited reactivity with species-specific DNA FISH probes and antibodies. **b.** Formalin-fixed, paraffin-embedded sections of human brain, liver, and kidney tissues were simultaneously stained with an antibody against Annexin V and a panel of species-specific DNA FISH probes and antibodies. Annexin V-positive apoptotic cells, but not Annexin V-negative viable cells, react with species-specific DNA FISH probes and antibodies. All experiments presented in Fig. 2 were performed twice by two different investigators.

In normal human tissues, formalin-fixed, paraffin-embedded sections of brain, liver, and kidney were stained with Annexin V antibody to identify the spontaneously occurring apoptotic cells among the viable cells. We found that in each case, only Annexin V-positive apoptotic cells reacted with FISH probes targeting DNA from pig, mouse, fungal, monkey, archaea, and fish species. In contrast, Annexin V-negative live cells were non-reactive (Fig. 2b, top row in each panel). Likewise, the Annexin V-positive apoptotic cells within these tissues reacted with antibodies against the Drosophila Wg protein, rat RTA1 protein, rice extensin glycoprotein, mouse MHC-II protein, algae CHR2 protein, and pig surfactant protein D. In contrast, the Annexin V-negative cells were non-reactive (Fig. 2b, bottom row in each panel).

### Phylogenetic diversity of ncDNA

We next tested the hypothesis that ncDNA represents a conglomeration of DNA– protein complexes from different species that become detectable following apoptosis. We had previously used a human long non-coding RNA probe comprising a set of Stellaris RNA FISH probes, each containing up to 48 unique sequences labelled with a fluorophore. We reported that the probe collectively bound to the non-coding RNA target transcript^2^. We found that, when tested against human DNA, the RNA probe aligned with the entire length of DNA with almost 100% coverage^2^. Since the non-coding RNA probe could also detect ncDNA, we used it to identify ncDNA in apoptotic bodies of hypoxia-induced apoptotic A375 cells. To generate apoptotic bodies, 1×10^6^ A375 cells were subjected to hypoxia-induced apoptosis, and 100 µL of the cell suspension containing approximately 1.5×10^5^ apoptotic cells was dropped onto glass slides from a height of 2 ft. This manoeuvre allowed the apoptotic cells to disperse into apoptotic bodies, which spread evenly on the glass slides. The slides were simultaneously probed with an antibody against Annexin V, a human whole-genome probe, and the non-coding RNA probe. The Annexin V-positive apoptotic bodies were found to co-localise with both the whole genomic and the non-coding RNA probes. Upon merging the images of whole genomic and non-coding RNA probe, 96.3% of the signals were found to co-localise, confirming that the non-coding RNA probe reliably detected ncDNA in virtually all apoptotic bodies (Fig. 3a). Consequently, all subsequent images obtained using the non-coding RNA probe were considered to represent ncDNA.

**Fig. 3:**
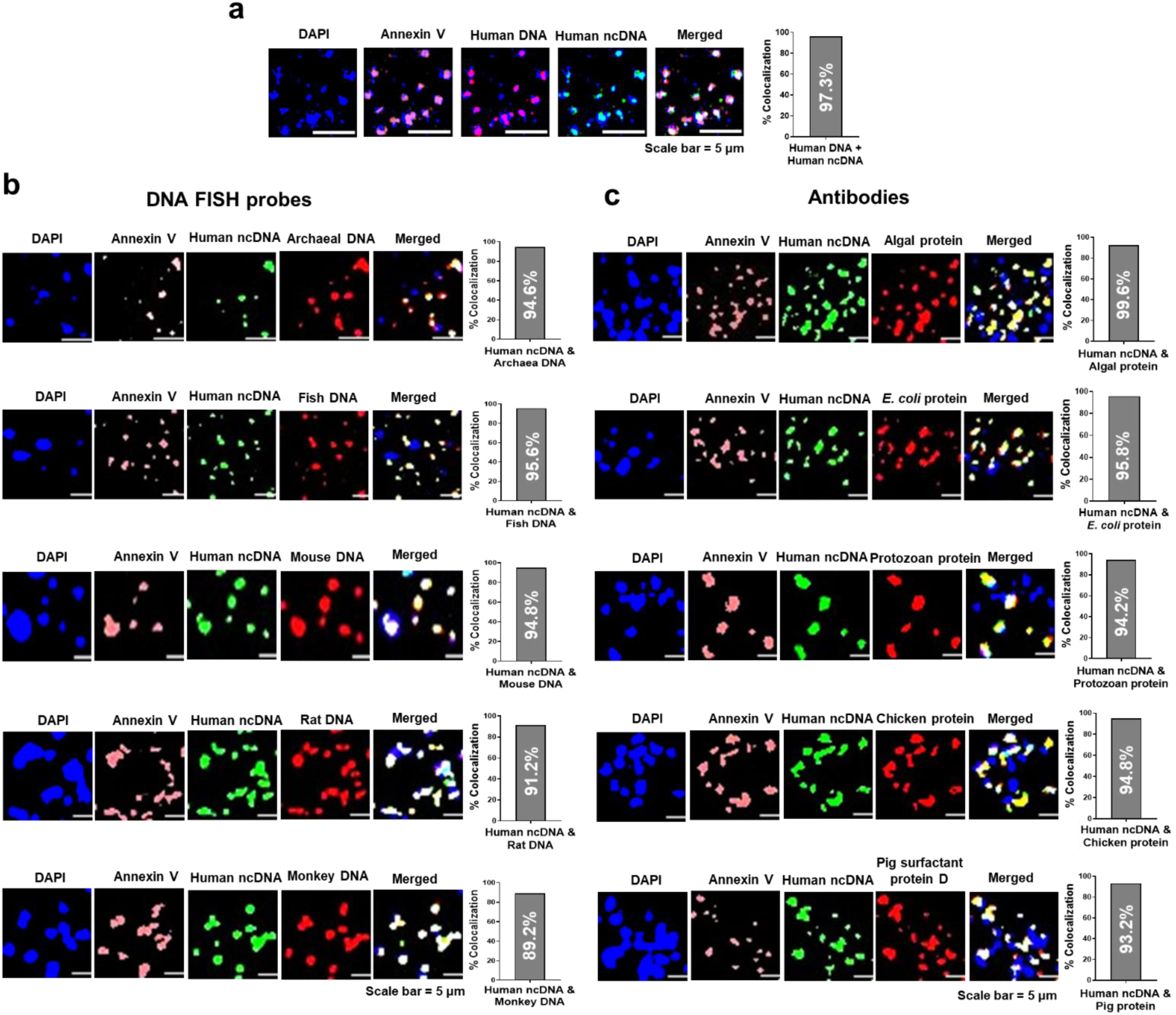
ncDNA in apoptotic bodies of A375 human melanoma cells reveals extensive phylogenetic diversity. **a.** Apoptotic bodies were simultaneously probed with an antibody against Annexin V (pink), a whole-genome DNA FISH probe (red), and a long non-coding RNA probe (green) to detect ncDNA. One thousand apoptotic bodies positive for human whole-genome DNA were analysed to determine ncDNA positivity, the percentage of apoptotic bodies that were also positive for ncDNA (histogram). A high degree of co-localisation of whole human genomic DNA and ncDNA (96.3%) is observed, indicating that nearly all apoptotic bodies were composed of ncDNA. The merged images exclude Annexin V signals. **b.** Apoptotic bodies were simultaneously probed with an antibody against Annexin V (pink), a long non-coding RNA probe (green), and a variety of species-specific DNA FISH probes (red). One thousand ncDNA-positive apoptotic bodies were analysed to determine the percentage of apoptotic bodies positive for species- specific DNA (histogram). A high degree of co-localisation of ncDNA and species- specific DNA is observed. For fluorescence colour consistency, FISH probes against pig, eubacteria, and fungi were pseudo-coloured as red, and the corresponding human long non-coding RNA probe was pseudo-coloured as green. Not all DAPI-stained apoptotic particles reacted with the Annexin V antibody, likely owing to uneven staining. The merged images exclude Annexin V signals. **c.** Apoptotic bodies were simultaneously probed with an antibody against Annexin V (pink), a long non-coding RNA probe (green), and various species-specific antibodies (red). One thousand apoptotic bodies positive for ncDNA were analysed to determine the percentage of apoptotic bodies positive for species-specific antibodies (histogram). A high degree of co-localisation of ncDNA and species- specific antibodies is observed. Not all DAPI-stained apoptotic particles reacted with Annexin V, likely owing to uneven staining of the apoptotic cells from which they were derived. The merged images exclude Annexin V signals. All experiments presented in Fig. 3 were performed twice by two different investigators.

We examined the apoptotic bodies of A375 cells using dual FISH probes against ncDNA and those against archaea, fish, mouse, rat, and monkey DNA (Fig. 3b). We found a high degree of co-localisation in each case (89.2–95.6%) between ncDNA and species-specific probes, as indicated in the adjacent histograms. We then performed immunoFISH using a FISH probe against ncDNA and antibodies against various species-specific proteins. We observed a similarly high degree of co-localisation of ncDNA and multiple proteins (range between 93.2% and 99.6%), as indicated in the adjacent histograms (Fig. 3c). These findings were further validated in human embryonic kidney (HEK-293) and human cervical cancer (HeLa) cells (Supplementary Fig. 3). Collectively, these results indicated that almost all apoptotic bodies derived from human cells consist of ncDNA containing sequences from various species, capable of synthesising their respective species-specific proteins that become detectable following apoptosis.

## Discussion

The present report is founded on two of our earlier studies^2,3^. The first study revealed that circulating cfChPs, mainly composed of ncDNA, that originate from apoptotic cells during normal cellular turnover can horizontally transfer to healthy cells. Once inside their new hosts, they randomly assemble into complex concatemers, some of which are ostensibly multi-megabase pairs in size, to become detectable by specific FISH probes and antibodies^2^. The second study indicated that high-molecular-weight DNA, when sonicated to generate 500–3000- bp fragments, similar in size to circulating cfChPs, can undergo unrestricted horizontal transfer into heterologous cells without being restricted by inter-specific and inter-kingdom boundaries^3^. Thus, sonicated particles from humans, bacteria, and plants could readily enter into nuclei of mouse cells, whereas those from humans, bacteria, and plants could spontaneously enter into bacteria. This observation begged the hypothesis that, over the course of evolution, DNA–protein complexes from dying organisms, especially during periods of severe environmental stress^4–6^, may have exchanged themselves between species and across kingdom boundaries, accumulated in recipient genomes and formed the dense patchwork of sequences that now constitute ncDNA. Findings from our first study suggested that components of these patchwork sequences become detectable using species-specific FISH probes and antibodies following cellular apoptosis. Our current results support the above hypothesis by demonstrating that

human cell apoptosis reveals the origin and nature of DNA–protein complexes that have accumulated over evolutionary time and are sequestered within human ncDNA. This implies that the phylogenetic diversity of ncDNA in every human cell represents a record of evolutionary history, reflecting a persistent species-spanning legacy. These findings lead us to propose that the silent ncDNA within human cells represents the remnants of an ancient genome once functional within primitive organisms, and that the 1% of protein-coding DNA gradually emerged to serve the complex functional requirements of higher-order organisms. Our earlier work^2^ had suggested that the ancient genome retains the potential to perform the various functions attributed to the modern genome, including DNA, RNA, and protein synthesis, guided by distinct biological rules that remain to be elucidated.

We were unable to confirm the phylogenetic diversity detected in the apoptotic cells using current DNA sequencing technologies, as the extensively fragmented apoptotic DNA failed quality control analysis, rendering it unfit for DNA sequencing.

Collectively, we propose that evolution may not have proceeded through random mutations in a hypothetical primordial genome, but rather through the progressive accumulation of DNA–protein complexes derived from diverse species, creating an ever-expanding genome that is now represented by ncDNA. In this framework, an appropriate combination of DNA–protein sequences that occurred by chance could have favoured the emergence of new species during the long course of evolution.

## Methods

### Ethics

This study was approved by the Institutional Ethics Committee (IEC) of the Advanced Centre for Treatment, Research, and Education in Cancer (ACTREC), Tata Memorial Centre (TMC), for the use of archival samples of normal human brain, liver, and kidney tissues (approval no. 900927).

### FISH probes and antibodies

A list of the FISH probes and antibodies used in this study, along with their species origin and procurement sources, is provided in Supplementary Table 1a–c. A list of the apoptosis detection reagents is provided in Supplementary Table 1d. FISH probes specific for human, mouse, pig, rat, and monkey were custom-synthesised, and their species specificity was certified by the respective companies (company names and their locations are provided in Supplementary Table 1a). DNA probes for eubacteria, fish, archaea, dogs, and fungi were procured commercially, and species specificity was certified by the respective manufacturers (the names of the companies and their country locations are listed in Supplementary Table 1a). Nonetheless, we independently tested each DNA FISH probe and antibody against cells and tissues of the various species examined to confirm species specificity and exclude cross-reactivity (Supplementary Fig. 1).

The human long non-coding RNA probe, composed of a set of Stellaris RNA FISH probes containing up to 48 unique sequences, each labelled with a fluorophore that collectively binds along an RNA target transcript, was procured commercially as indicated. For fluorescent colour compatibility, two differently labelled long non-coding RNA probes (red and green) were used as required.

### Cell lines and culture

A list of cell lines used in this study, their tissue origins, culture media used, and procurement sources is provided in Supplementary Table 2. The cells were cultured in 35-mm dishes at 37 °C, 95% air, 5% CO2, and 95% humidity. For the experiments, 6×10^4^ cells were plated per dish and used the following day when the cell density was approximately 1×10^5^.

### Flow cytometry

Cells were incubated in a hypoxic chamber (1% O2) for 72 h, harvested by gentle scraping, pelleted, and resuspended in Annexin binding buffer. To identify apoptotic cells, cells were stained with FITC-labelled Annexin V for 20 min in the dark and sorted using a FACS Aria III (BD Biosciences, San Jose, CA, USA) with FACS Diva software (version 4.0.1.2; Becton, Dickinson and Company). Cells were gated based on forward- and side-scatter characteristics, and Annexin V-positive cells were sorted, collected in PBS, and processed for IF and FISH.

### IF and FISH

#### IF and FISH on cultured cells

The various cell lines were cultured in 35-mm dishes in Dulbecco’s modified Eagle’s medium (DMEM) supplemented with 10% foetal bovine serum (FBS) in a humidified incubator at 37 °C with 5% CO2. Cells (1×10^5^) were harvested, cytospun onto positively charged slides, and fixed in 4% paraformaldehyde (PFA). For IF analysis, the unbound PFA was neutralised with 0.3 M glycine, treated with saponin buffer containing 5% normal goat serum (1 h), and immunostained with appropriate primary and secondary antibodies. The slides were mounted using VECTASHIELD DAPI (Vector Laboratories, Newark, CA, USA, cat. no. H1200). For the FISH analysis, the cells were dehydrated in a series of molecular-grade ethanol (70%, 80%, and 100%). They hybridized with the appropriate FISH probes diluted in hybridization buffer (10% dextran sulfate, 10% deionized formamide, and 2X SSC dissolved in nuclease-free water) in a humidified chamber at 37°C overnight (16 hours). Slides were washed in 2×SSC for 15 min at 37°C and in 1×SSC at room temperature (RT) for 10 min. The slides were mounted using VECTASHIELD DAPI. For both IF and FISH, the images were acquired using an Applied Spectral Bio-Imaging System (Yokne’am Illit, Israel).

#### IF and FISH on formalin-fixed paraffin-embedded tissue sections

Formalin-fixed paraffin-embedded sections (5 µm thickness) of human brain, liver, and kidney were deparaffinised thrice in xylene (100%), re-hydrated through an alcohol series (100% thrice, 80%, 70% and 50%), and distilled water, and subjected to antigen retrieval at 90 °C using sodium citrate buffer (pH 6) for 20 min. Slides were cooled to RT (approximately 25–30 °C), treated with saponin buffer (1 h), and stained with Annexin V. Slides were then fixed in 2% PFA for 10 min and subjected to FISH as described in the previous section.

#### IF and FISH on apoptotic bodies

To generate apoptotic bodies, 1×10^6^ A375 melanoma cells were subjected to hypoxia-induced apoptosis. The apoptotic cell suspension containing approximately 1.5×10^5^ apoptotic bodies was diluted to 100 µL phenol red-free DMEM and dropped onto glass slides from a height of 2 ft to evenly disperse the apoptotic bodies. Slides were then processed for IF and FISH (described in the subsequent section).

### Attempts to sequence apoptotic DNA

We extracted DNA from apoptotic A375 human melanoma cells to confirm our FISH findings of phylogenetic diversity using whole-genome sequencing.

However, DNA from two biological replicates failed quality assessment, which precluded genomic sequencing. One of the two agarose gel images is provided in Supplementary Fig. 4.

### Image acquisition and processing

For fluorescence microscopy, we utilized the Spectral Bio-Imaging System, which is installed on a BX63 Olympus microscope. Images were captured under a 40× air objective lens using FISH View Software 8.0. All images were captured at identical optical settings for comparison and to prevent saturation and photo bleaching. All images were processed using FISH View Software 8.0.

### Use of a long non-coding RNA probe to detect ncDNA

We previously used a human long non-coding RNA probe comprising a set of Stellaris RNA FISH probes, each containing a pool of up to 48 unique sequences labelled with a fluorophore^2^. The probe collectively binds to the non-coding RNA target transcript^2^. We reported that, when tested against human DNA, the RNA probe aligned with the full DNA length with almost 100% coverage^2^. Leveraging the ability of the non-coding RNA probe to detect ncDNA, we used it to identify ncDNA. Apoptotic bodies were simultaneously probed with antibodies against Annexin V, the non-coding RNA probe, and various species-specific FISH probes.

### Procurement and processing of positive-control cells and tissues

Sources of cells and tissues used as positive controls are provided in Supplementary Table 3. Cells and tissues were processed as follows: archaebacteria, eubacteria (E. coli), fungus (yeast), and protozoa (Entamoeba histolytica) cells were fixed in 4% PFA, smeared onto slides, air-dried, and processed for FISH and IF. Algal samples (moss tissue) were washed with PBS, fixed in 4% PFA, smeared on slides, air-dried, and processed for FISH and IF. NIH3T3 mouse fibroblasts, dog cells, and monkey cells were cytospun onto slides, fixed in 4% PFA, and processed for FISH and IF. Drosophila lymph gland, chicken liver, rat liver, and pig liver tissues were fixed in 10% neutral-buffered formalin, embedded in paraffin blocks, and sectioned for FISH and IF.

### Attempts to sequence apoptotic DNA

We extracted DNA from apoptotic A-375 human melanoma cells to confirm our FISH findings of phylogenetic diversity in apoptotic cells using whole-genome sequencing. However, the DNA from two biological replicates failed the quality check, which precluded any genomic sequencing.

### Statistical analyses

Our study did not involve any formal statistical analysis.

### Data availability

All the data are included in the manuscript. Additional data will be provided upon reasonable request to the corresponding author.

## Acknowledgements

We thank Mr. Ashish Pawar for his help in preparing the manuscript.

## Author contributions

R.J., A.R., L.K., R.S., M.M. and S.B. performed the immunofluorescence and FISH experiments. R.L. was responsible for cell culture and provided the live and hypoxia-induced apoptotic cells. G.V.R. supervised the immunofluorescence and FISH experiments and wrote the manuscript. S.S. performed flow cytometry-based sorting experiments, supervised the IF and FISH analyses and wrote the manuscript. I.M. conceptualised and was responsible for overall supervision of the work, procured funding, wrote the manuscript and finalised and approved the final version.

## Competing interests

The authors declare no conflict of interest.

## Materials & Correspondence

Correspondence and requests for materials should be addressed to Prof. Indraneel Mittra.

Reprints and permissions information is available at www.nature.com/reprints.

## Additional information

Supplementary Information is available for this paper.

## Supplementary Information

### Supplementary figures

**Supplementary Fig. 1.**
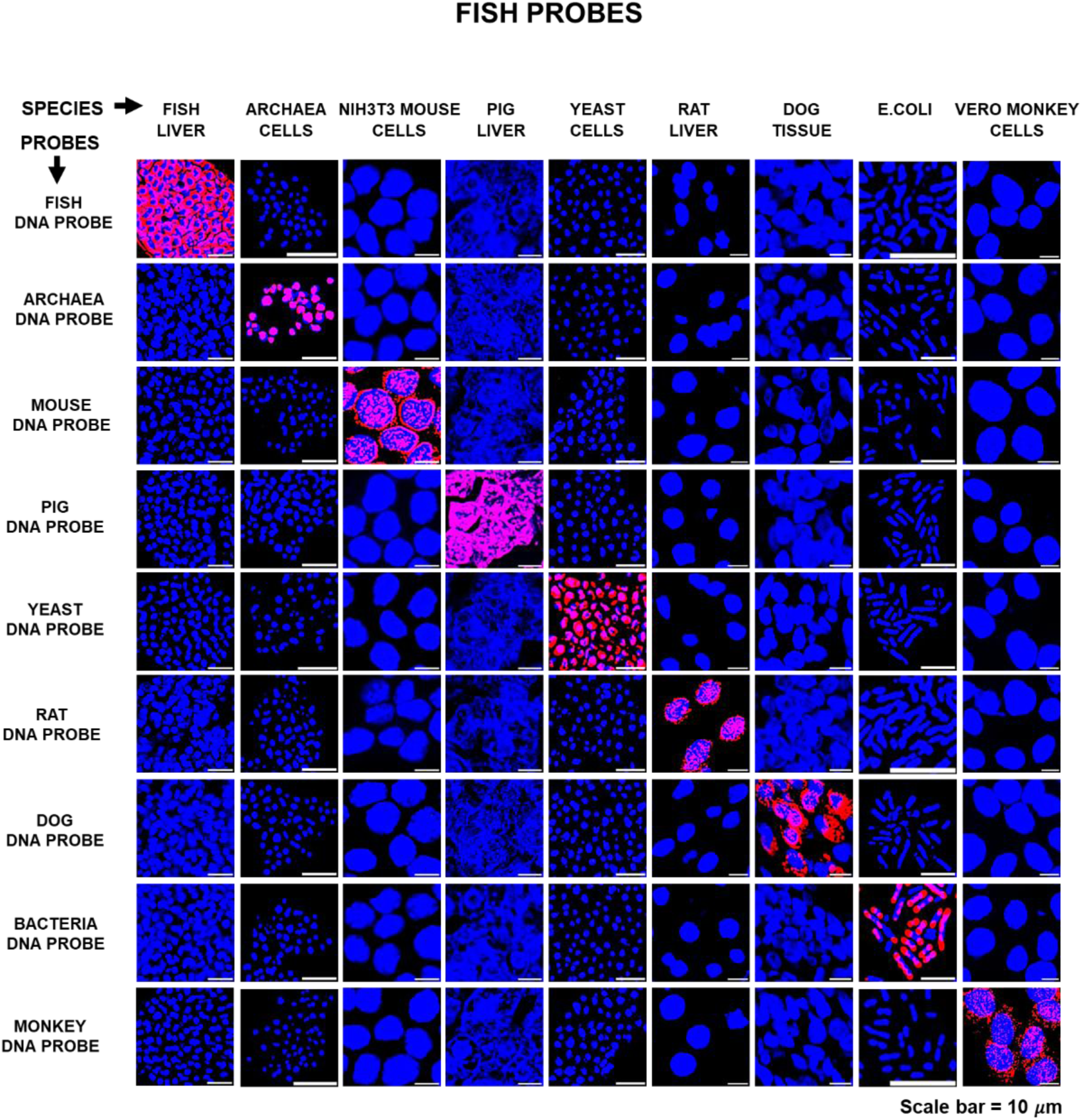
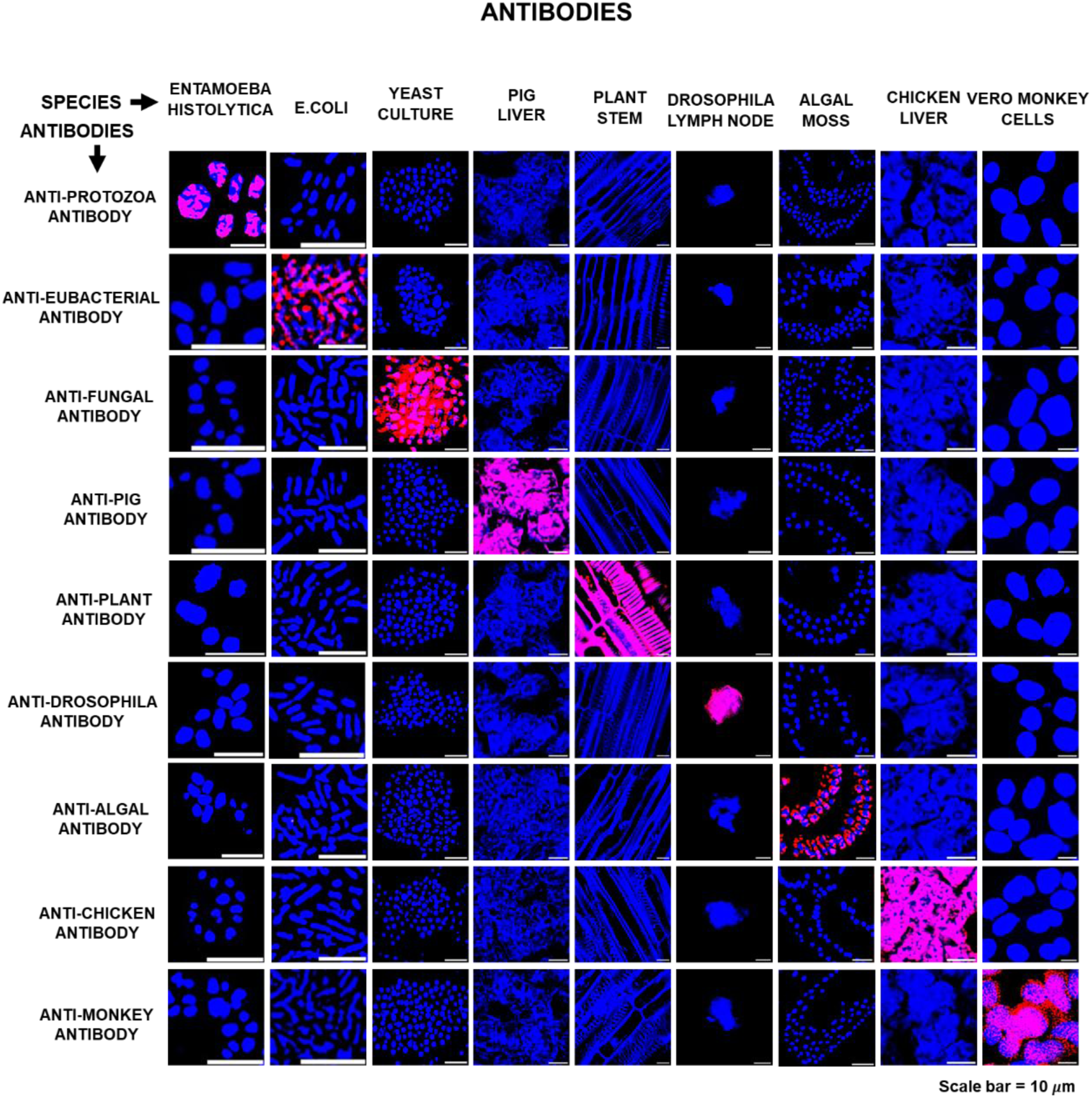
Experiments to confirm species-specificity of FISH probes and antibodies. **a.** Cells were fixed with paraformaldehyde for 10 min, whereas formalin-fixed, paraffin-embedded sections were used in the case of tissues. Cells and tissue sections were probed with species-specific probes as indicated. **b.** Same as in (a), except cells and tissue sections were probed with appropriate species-specific antibodies as indicated.

**Supplementary Fig. 2.**
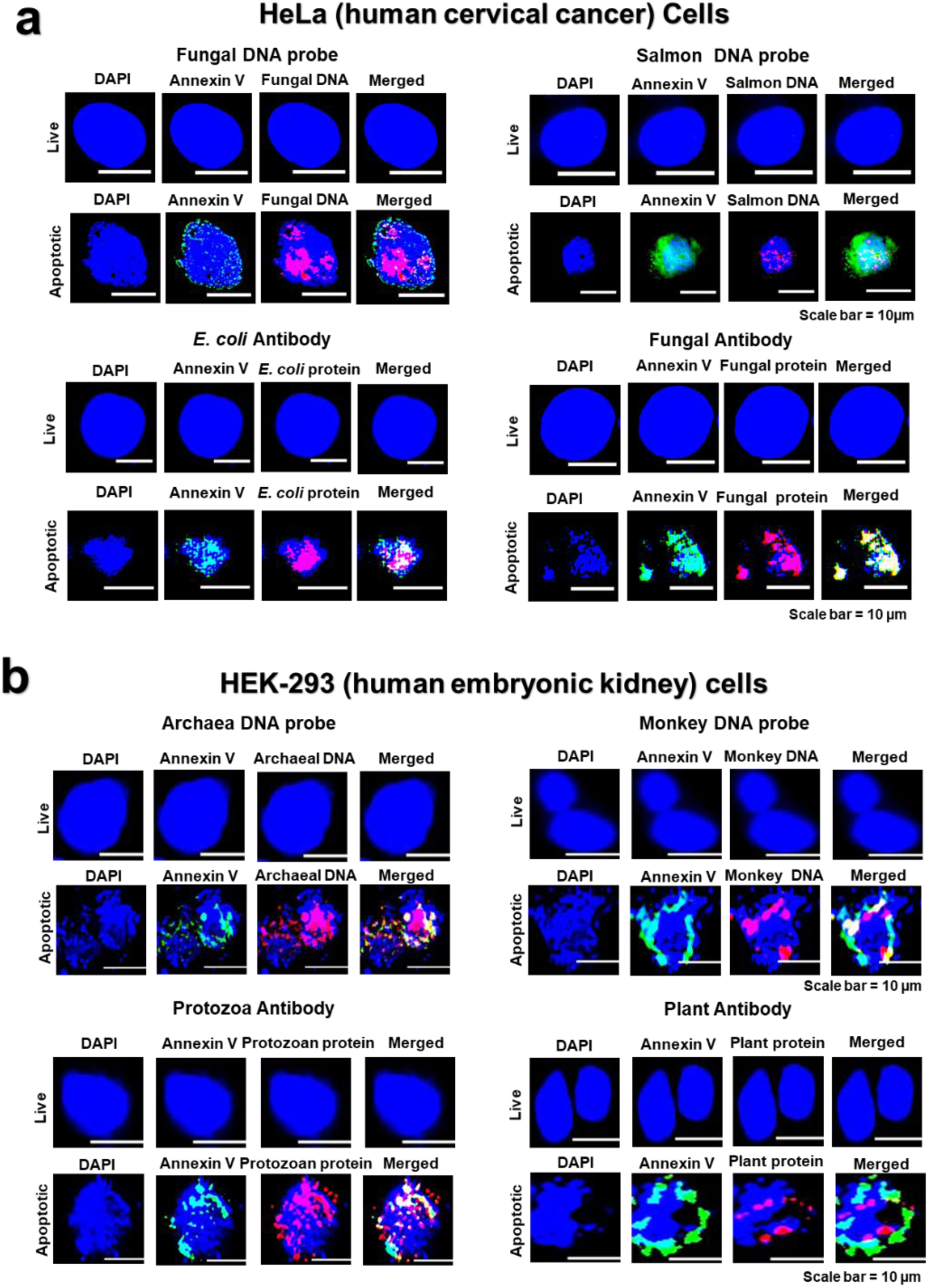
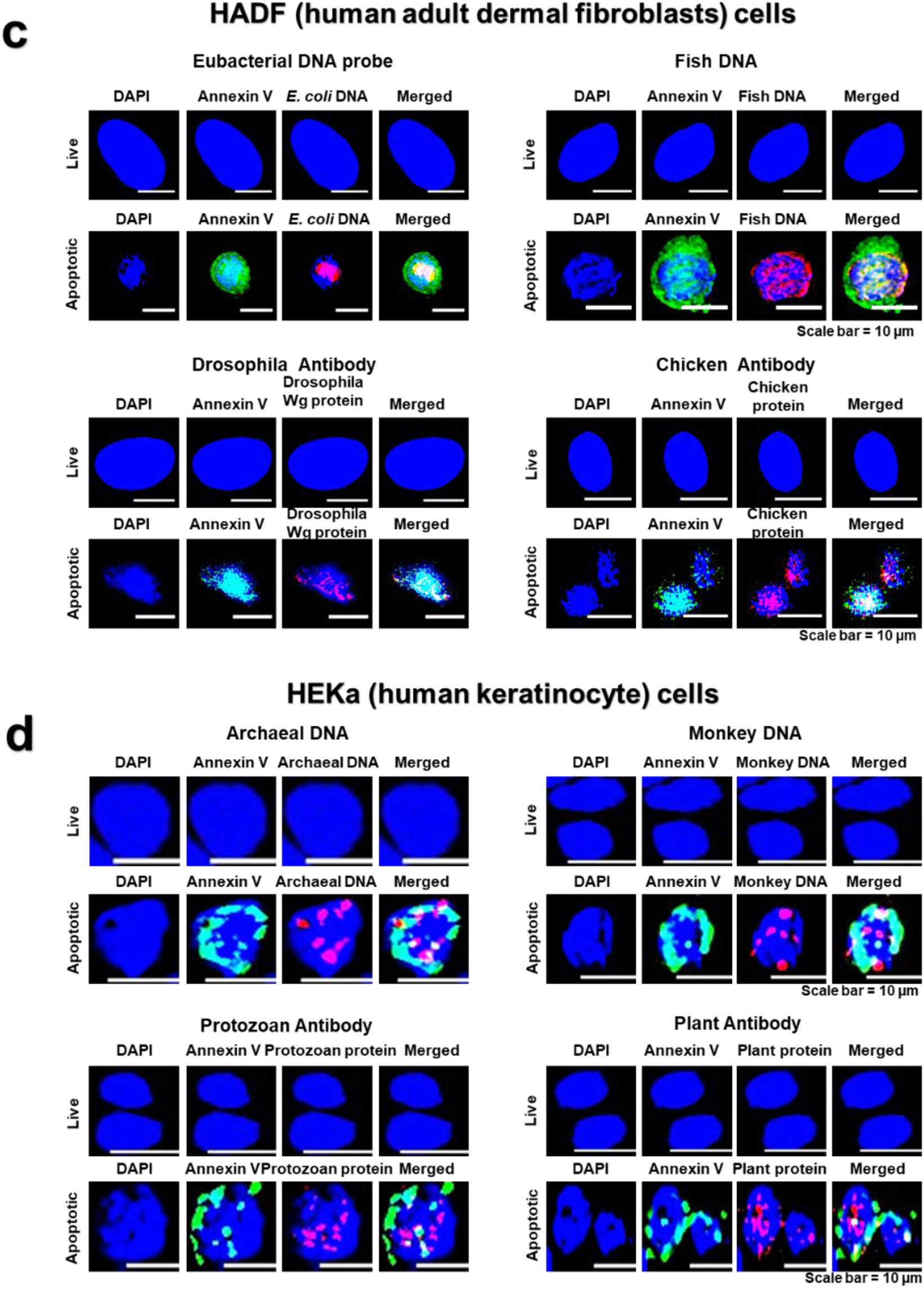
Apoptosis of diverse human cell types reveals their phylogenetic diversity. **a–d.** HeLa (human cervical cancer), HEK-293 (human embryonic kidney), human dermal fibroblast cells, and human keratinocytes. Methodological details are provided in Fig. 1.

**Supplementary Fig. 3:**
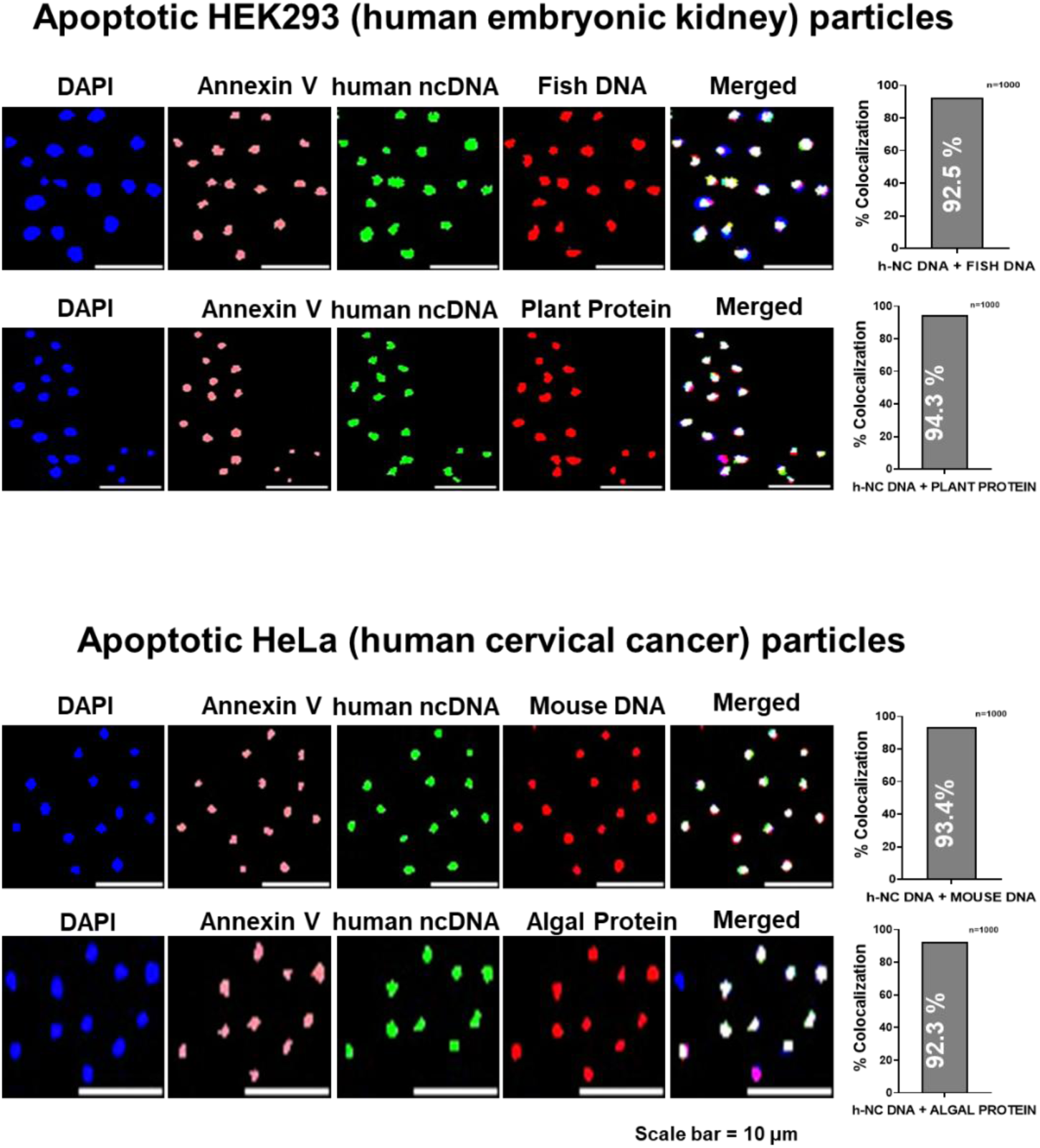
Non-coding DNA in apoptotic bodies of cell types other than A375 human melanoma cells reveals their phylogenetic diversity. Methodological details are provided in Fig. 3.

**Supplementary Fig. 4:**
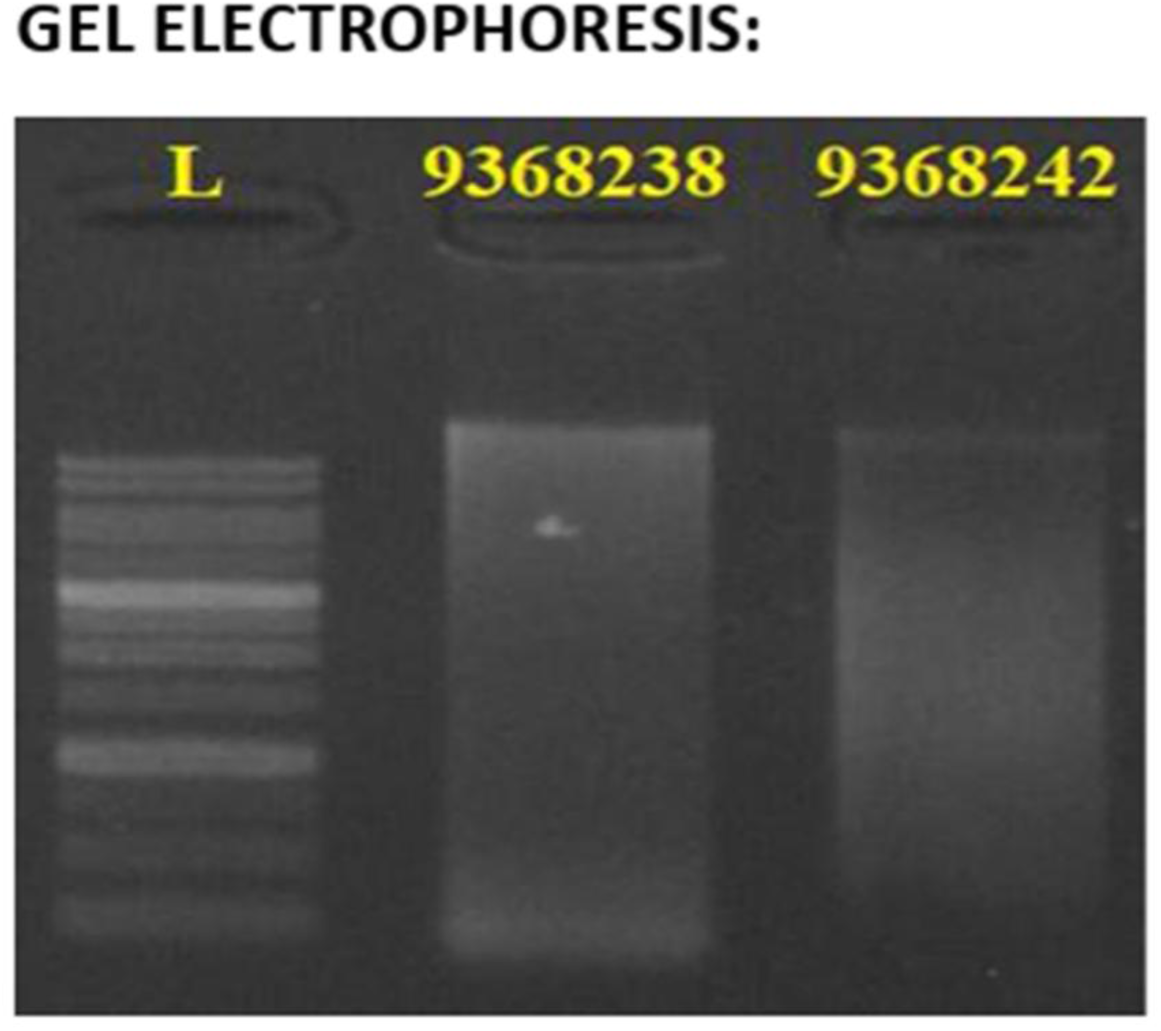
Agarose gel electrophoresis of hypoxia-induced apoptotic cells showing a smear pattern, which failed the quality-control test for whole genome sequencing.

### Supplementary tables

**Supplementary Table 1a.**
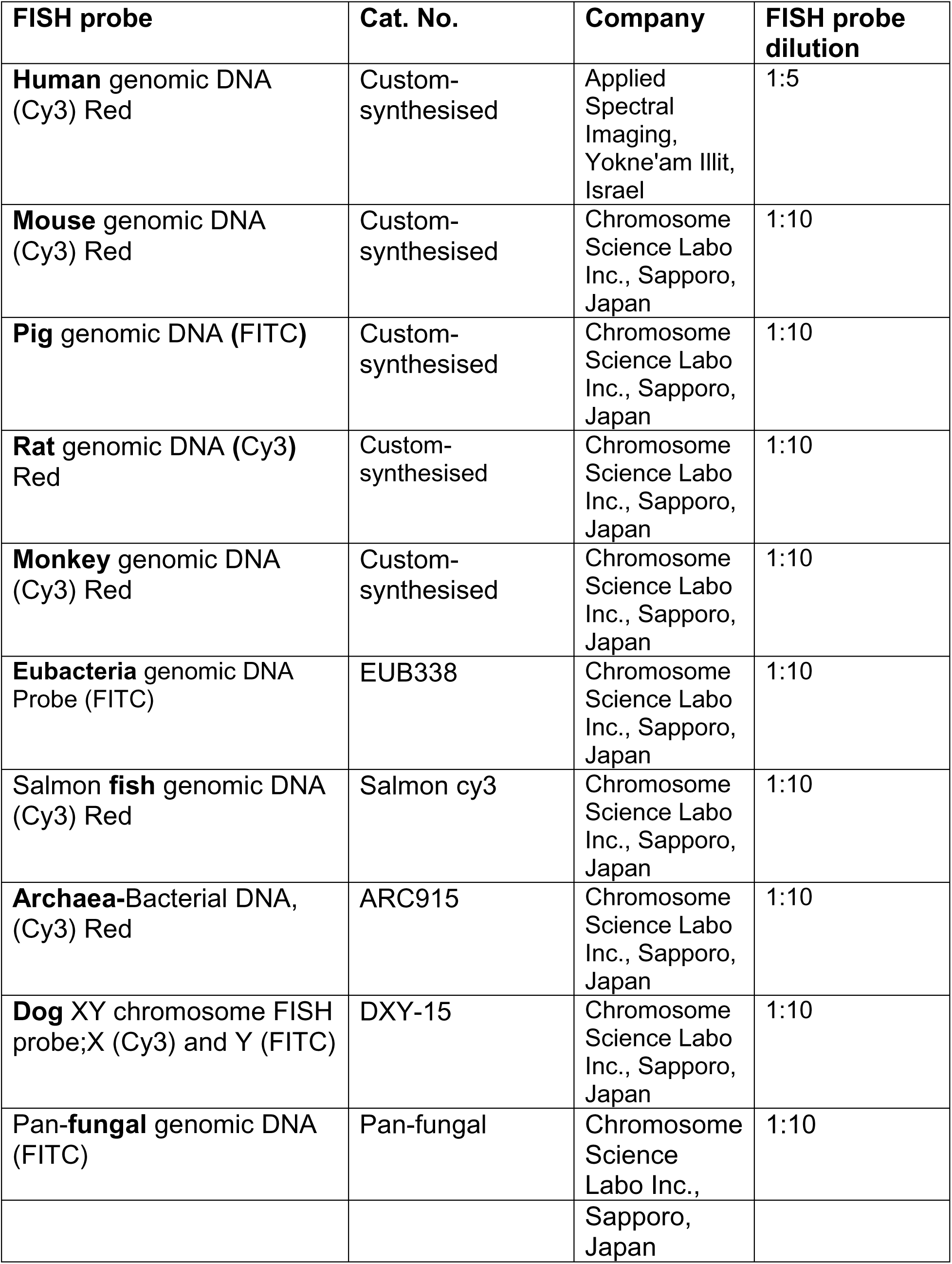
DNA FISH probes.

| **FISH probe** | **Cat. No.** | **Company** | **FISH probe dilution** |
| --- | --- | --- | --- |
| **Human** genomic DNA (Cy3) Red | Custom-synthesised | Applied Spectral Imaging, Yokne'am Illit, Israel | 1:5 |
| **Mouse** genomic DNA (Cy3) Red | Custom-synthesised | Chromosome Science Labo Inc., Sapporo, Japan | 1:10 |
| **Pig** genomic DNA (FITC) | Custom-synthesised | Chromosome Science Labo Inc., Sapporo, Japan | 1:10 |
| **Rat** genomic DNA (Cy3) Red | Custom-synthesised | Chromosome Science Labo Inc., Sapporo, Japan | 1:10 |
| **Monkey** genomic DNA (Cy3) Red | Custom-synthesised | Chromosome Science Labo Inc., Sapporo, Japan | 1:10 |
| **Eubacteria** genomic DNA Probe (FITC) | EUB338 | Chromosome Science Labo Inc., Sapporo, Japan | 1:10 |
| Salmon **fish** genomic DNA (Cy3) Red | Salmon cy3 | Chromosome Science Labo Inc., Sapporo, Japan | 1:10 |
| **Archaea**-Bacterial DNA, (Cy3) Red | ARC915 | Chromosome Science Labo Inc., Sapporo, Japan | 1:10 |
| **Dog** XY chromosome FISH probe;X (Cy3) and Y (FITC) | DXY-15 | Chromosome Science Labo Inc., Sapporo, Japan | 1:10 |
| **Pan-fungal** genomic DNA (FITC) | Pan-fungal | Chromosome Science Labo Inc., | 1:10 |
|  |  | Sapporo,<br>Japan |  |

**Supplementary Table 1b.**
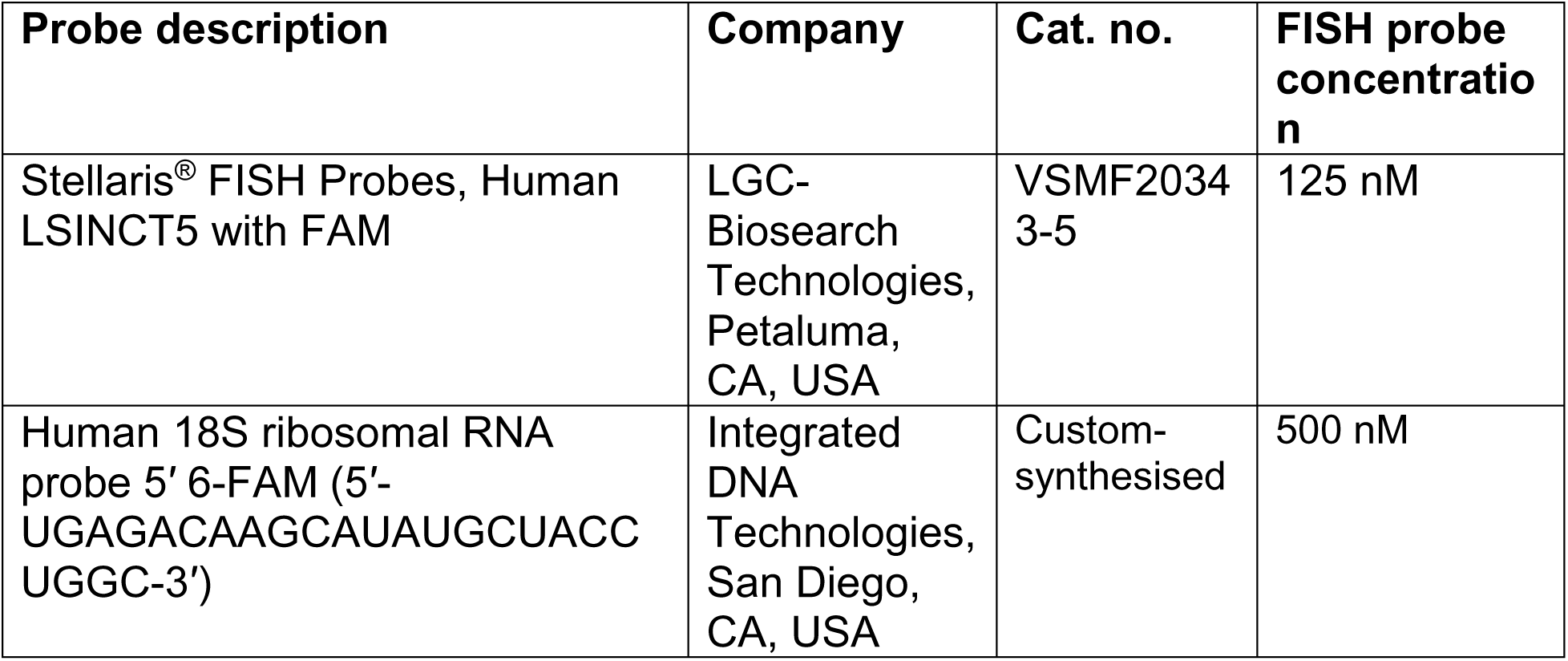
Long non-coding RNA probes.

**Supplementary Table 1c.**
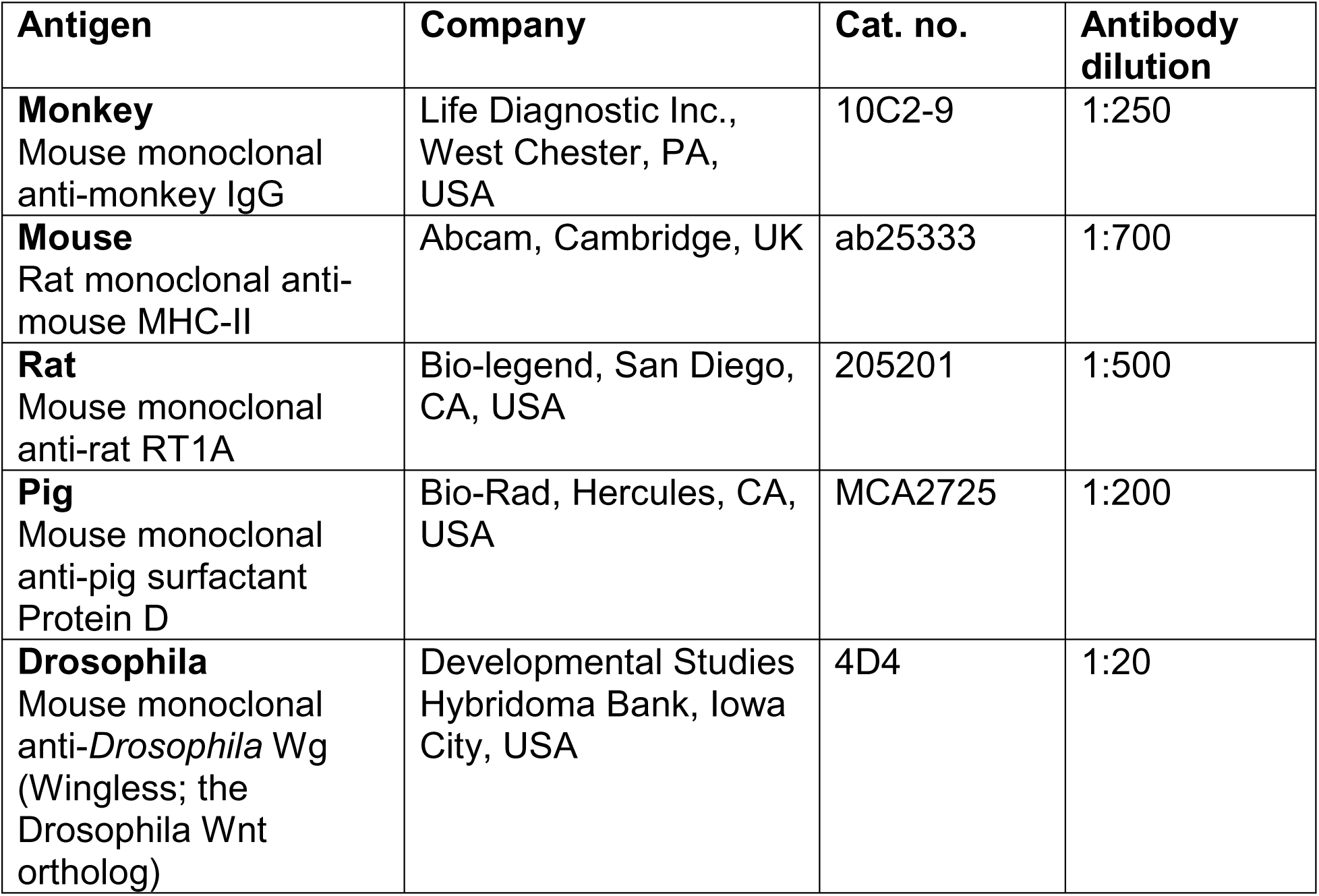
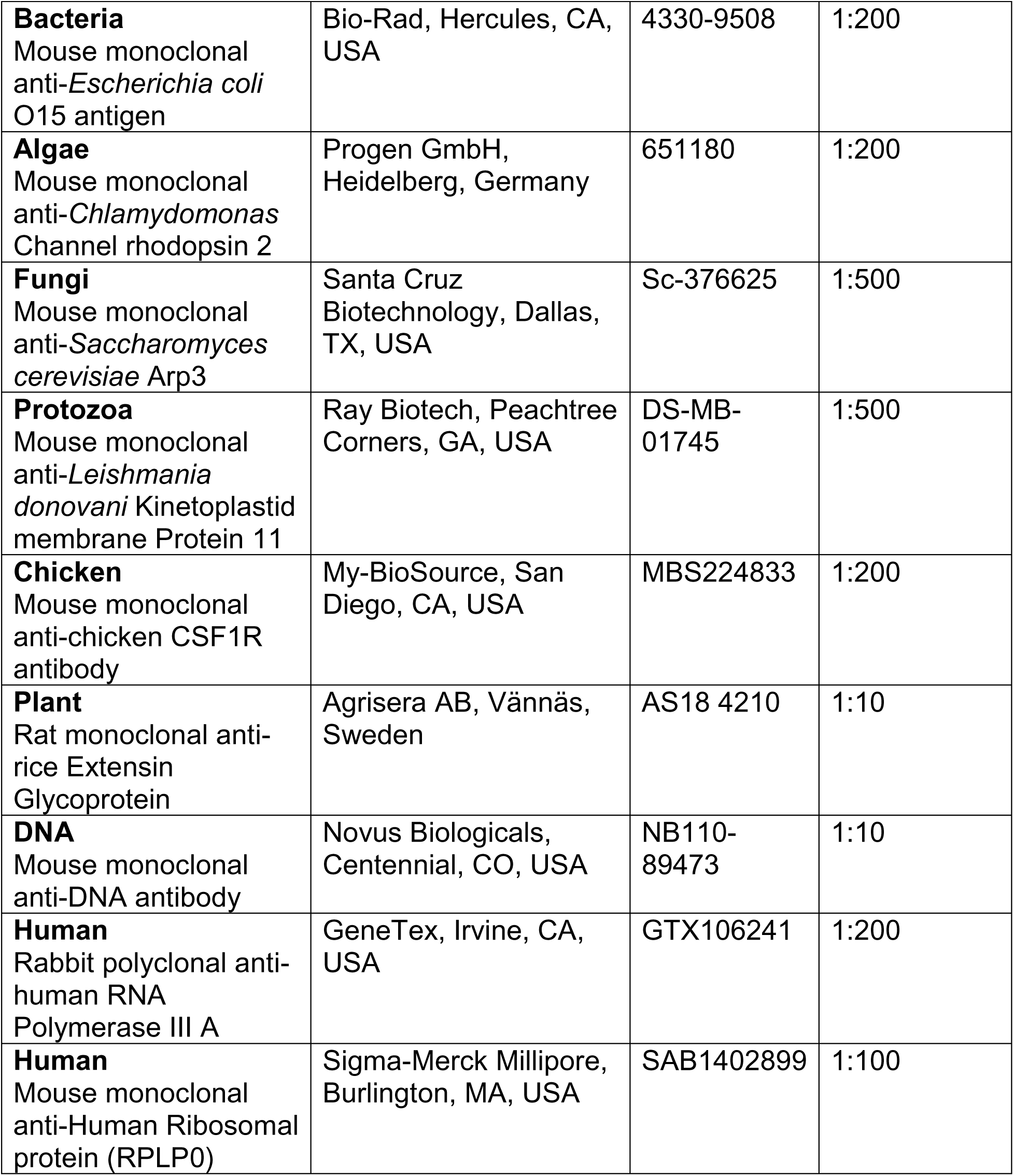
Antibodies.

**Supplementary Table 1d.** Apoptosis detection reagents.

| Reagent name | Company | Cat. no. | Reagent dilution |
| --- | --- | --- | --- |
| BD Pharmingen™ FITC Annexin V Apoptosis Detection Kit | BD Biosciences, San Jose, CA, USA | 556547 | 1:20 |
| BD Pharmingen™ PE Annexin V Apoptosis Detection Kit | BD Biosciences, San Jose, CA, USA | 559763 | 1:20 |
| Rabbit polyclonal anti-Annexin V antibody | Antibodies.com, Cambridge, UK | A13619 | 1:100 |
| Goat polyclonal anti-rabbit IgG (H+L) secondary antibody, DyLight-633 | ThermoFisher Scientific, Waltham, MA, USA | 35562 | 1:500 |

**Supplementary Table 2.** List of cell lines.

| Cell line | Tissue Origin | Source | Culture medium used |
| --- | --- | --- | --- |
| A375 | Human melanoma | ATCC (CRL-1619) | DMEM + 10% FBS |
| MRC-5 | Human lung fibroblast | ATCC (CCL-212) | EMEM + 10% FBS |
| MDA-MB-231 | Human breast cancer | ATCC (HTB-26) | DMEM + 10% FBS |
| HEK-293 | Human embryonic kidney | ATCC (CRL-1573™) | DMEM + 10% FBS |
| HeLa | Human cervical cancer | ATCC (CRM-CCL-2™) | DMEM + 10% FBS |
| HADF | Human dermal fibroblast | HiMedia (CL005) | DMEM + 10% FBS |
| HEKa | Human keratinocyte | ThermoFisher Scientific (C0055C) | DMEM + 10% FBS |

DMEM, Dulbecco's Modified Eagle Medium; EMEM, Eagle's Minimum Essential Medium; FBS, Foetal bovine serum

**Supplementary Table 3.**
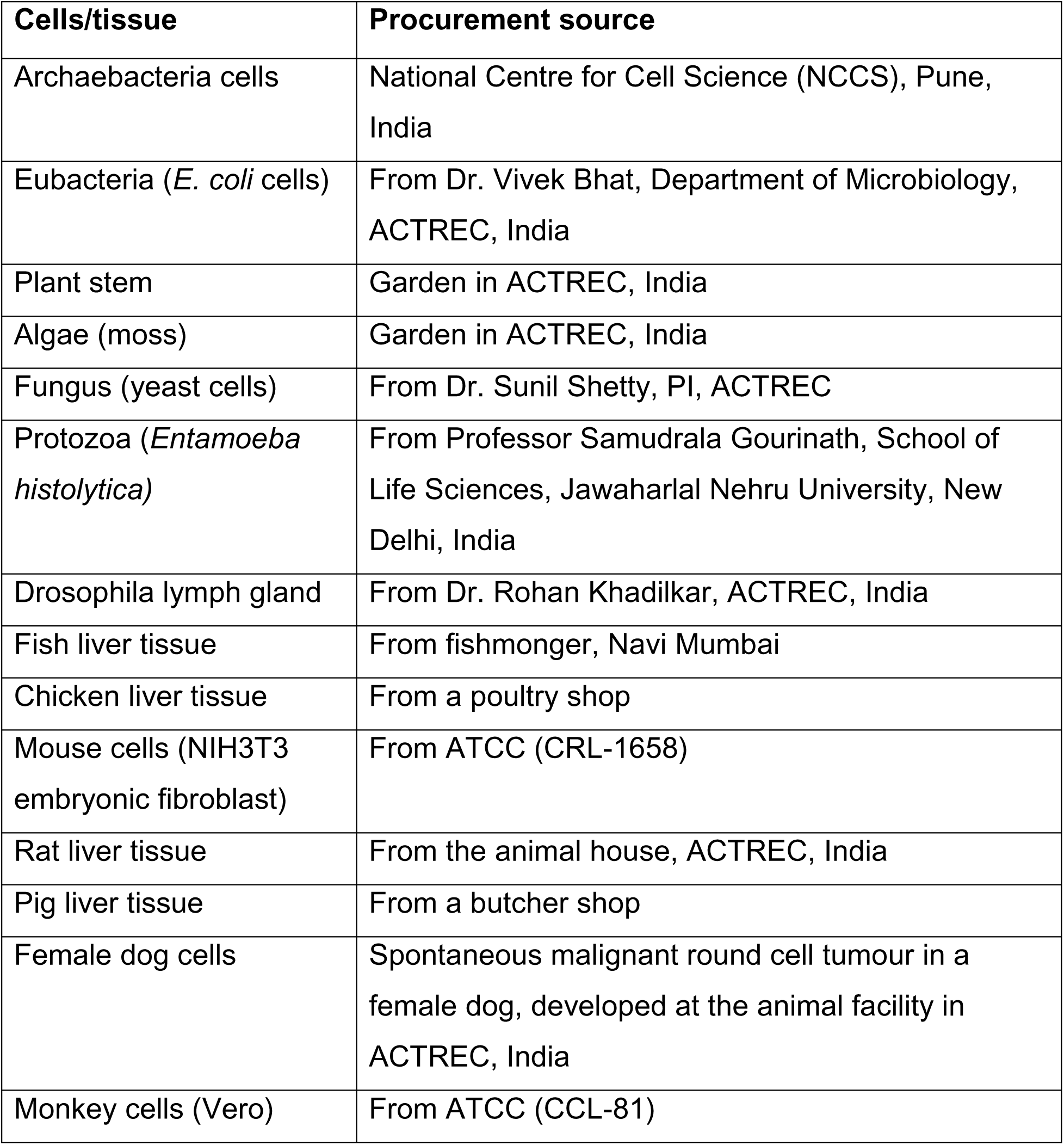
Positive-control cells and tissues.

| Cells/tissue | Procurement source |
| --- | --- |
| Archaeobacteria cells | National Centre for Cell Science (NCCS), Pune, India |
| Eubacteria (*E. coli* cells) | From Dr. Vivek Bhat, Department of Microbiology, ACTREC, India |
| Plant stem | Garden in ACTREC, India |
| Algae (moss) | Garden in ACTREC, India |
| Fungus (yeast cells) | From Dr. Sunil Shetty, PI, ACTREC |
| Protozoa (*Entamoeba histolytica*) | From Professor Samudrala Gourinath, School of Life Sciences, Jawaharlal Nehru University, New Delhi, India |
| Drosophila lymph gland | From Dr. Rohan Khadilkar, ACTREC, India |
| Fish liver tissue | From fishmonger, Navi Mumbai |
| Chicken liver tissue | From a poultry shop |
| Mouse cells (NIH3T3 embryonic fibroblast) | From ATCC (CRL-1658) |
| Rat liver tissue | From the animal house, ACTREC, India |
| Pig liver tissue | From a butcher shop |
| Female dog cells | Spontaneous malignant round cell tumour in a female dog, developed at the animal facility in ACTREC, India |
| Monkey cells (Vero) | From ATCC (CCL-81) |

## Notes

### Competing Interest Statement

The authors have declared no competing interest.

### Summary of Updates

1) We have modified the title 2) We have modified the abstract 3) We have modified the introduction section 4) We have modified the discussion section

### Funding

- Department of Atomic Energy: CTC-TMC

